# Behavioural and Physiological Signatures of Chronic Unpredictable Mild Stress Models in Mice

**DOI:** 10.64898/2025.11.30.691402

**Authors:** Ines Erkizia-Santamaría, Luna Fernández, Igor Horrillo, J. Javier Meana, Jorge E. Ortega

**Affiliations:** Department of Pharmacology, University of the Basque Country UPV/EHU, Leioa, Bizkaia, Spain; Institute of Pharmacology and Neurosciences, Universidade de Lisboa, Lisboa, Portugal; Centro de Investigación Biomédica en Red de Salud Mental, Instituto de Salud Carlos III, Spain; Biobizkaia Health Research Institute, Barakaldo, Bizkaia, Spain

**Keywords:** Chronic stress, Depression, Chronic Unpredictable Mild Stress, Depression-like Model, Anxiety

## Abstract

The chronic unpredictable mild stress (CUMS) paradigm is one of the most widely used preclinical models to investigate the role of stress in the neurobiology of depression and to evaluate potential antidepressant therapies, owing to its strong translational relevance. However, despite its extensive use, substantial discrepancies and concerns regarding the reproducibility of CUMS outcomes persist in the literature, with many studies reporting inconsistent or unreliable results. In the present comparative study, we implemented several CUMS protocols in mice that differed in the intensity and duration of stress exposure, and systematically evaluated the resulting pathophysiological and behavioural alterations associated with depressive- and anxiety-like phenotypes. We found that progressive escalation in stress intensity and frequency, as well as prolonged exposure, induced increasingly robust disease-related behavioural and physiological changes, including depression- and anxiety-like behaviours, adrenal hypertrophy, and reduced body weight gain.. Moreover, social isolation emerged as a major contributing factor exacerbating stress-induced deficits. Altogether, data suggest that a multidimensional assessment of the behavioural impairments is critical to reinforce the physiological changes indicative of a hyperactive hypothalamic-pituitary-adrenal axis (HPA). Differences in individual stress susceptibility, stressor intensity, frequency, and housing conditions are key determinants of CUMS outcomes. Therefore, precise, detailed, and transparent reporting of protocol parameters is critical to enhance reproducibility and enable meaningful comparisons across studies.

## 1. INTRODUCTION

Major depressive disorder (MDD), commonly referred to as depression, constitutes the second most prevalent mental disease worldwide, and is the most burdensome across all ages in the global population (GBD 2017 Disease and Injury Incidence and Prevalence Collaborators, 2018; GBD 2019 Mental Disorders Collaborators, 2022). Moreover, the prevalence of depressive disorders is increasing, according to trend analyses (Moreno-Agostino et al., 2021). MDD is characterized by core symptoms such as persistent low mood, diminished motivation, and loss of interest or pleasure, a condition known as anhedonia. Additional features may include cognitive deficits, sleep disturbances, and psychomotor retardation. These diverse manifestations contribute to the marked heterogeneity of MDD in both clinical presentation and disease course, which may be episodic, recurrent, or chronic (American Psychiatric Association, 2013; Gururajan et al., 2019). Furthermore, clinical anxiety constitutes the most prevalent comorbidity of MDD, experienced by over 60% of patients, with anxiety symptoms often arising 1-2 years before depressive symptoms (Demyttenaere & Heirman, 2020; Malhi & Mann, 2018).

The undefined etiopathogenesis of most psychiatric conditions, including MDD, is thought to underlie both the clinical heterogeneity observed among individuals with the same diagnosis and their differential responsiveness to pharmacological interventions (Nutt, 2025). Although depressive disorders are characterized by complex and heterogeneous biological underpinnings, including alterations in inflammatory pathways, dysregulation of the hypothalamic-pituitary-adrenal (HPA) axis, and imbalances in autonomic nervous system activity (Nedic Erjavec et al., 2021), several specific risk factors have been identified. Chronic stress is widely recognized as a key etiological factor in MDD, with exposure to significant life stressors representing one of the most robust predictors of depressive episode onset (Hill et al., 2012). Extensive evidence supports the link between chronic stress, HPA dysfunction and depressive disorders (Dwyer et al., 2020; Kessler, 1997). Prolonged exposure to elevated glucocorticoid levels induces dendritic retraction and neuronal atrophy in the frontal cortex (FC) and hippocampus (HC), regions that are critical for emotional regulation and cognitive processing. These alterations closely parallel the structural abnormalities reported in the brains of patients with depression (Duman & Aghajanian, 2012).

Both the biological features of stress neurobiology and the impairments that comprise a dysregulated stress system show remarkable translation from humans to rodents (Bale et al., 2019). Therefore, chronic stress models arguably have the best construct and face validity among rodent models developed for the study of depression-related disorders. The chronic unpredictable mild stress (CUMS) model, based on a series of observations made by Katz and colleagues in the early 1980s (Katz, 1982), has evolved significantly since it was first developed, and remains one of the most robust, reliable and widely used stress-based depression-like models (Planchez et al., 2019; Willner, 2017b). CUMS models recapitulate behavioural disturbances associated to depression and evoke an array of neurobiological changes that mirror those observed in depressive disorders (Hill et al., 2012; Strekalova et al., 2022). Despite the widespread use of CUMS models over several decades, major discrepancies and concerns regarding reproducibility persist in the literature. Numerous studies have reported unreliable and inconsistent outcomes, particularly in relation to the induction of anhedonia (Antoniuk et al., 2019; Strekalova et al., 2022). As a result, the interpretation of data derived from CUMS paradigms in the framework of depressive disorders remains problematic (Anisman & Matheson, 2005; Strekalova et al., 2022). In this context -marked by the urgent need for novel and more effective antidepressant treatments, including the emerging class of fast-acting compounds such as psychedelics-it is imperative to implement well-defined and standardized stress protocols capable of reliably eliciting disease-relevant phenotypic features. The development of such robust preclinical models is essential not only for enhancing translational validity, but also for ensuring the reliability of pharmacological screening efforts in this rapidly evolving field (Erkizia-Santamaria et al., 2025a).

Experimental protocols to generate CUMS models consist of chronic stress paradigms that combine diverse physical and psychosocial stressors of unpredictable nature, of variable frequency and for an undefined duration of several weeks, under varying housing conditions. In view of the high methodological inconsistencies found in the literature, there is a compelling need for protocol standardization, in the interest of enhancing the reliability and reproducibility of the model across laboratories. In the present comparative study, we implemented distinct CUMS protocols in mice, each differing in the intensity and/or duration of stress exposure, and critically evaluated the resulting impairments related to depressive-like or anxiety-like symptomatology and pathophysiological alterations. All protocols were conducted using the same mouse strain, experimental facilities, and were performed by the same personnel, which allowed for a more exhaustive and controlled comparison than that afforded by cross-laboratory studies. This strategy aimed to identify key variables contributing to the consistent and reliable induction of a CUMS-based depression-like phenotype.

## 2. MATERIALS AND METHODS

### 2.1 Animals

Adult male C57BL/6J mice (8 weeks old) were purchased from Envigo (Barcelona, Spain) and housed under standard laboratory conditions on a 12 h light/dark cycle, at room temperature (22–24 °C), with food and water available *ad libitum*. The animal care and experimental protocols were carried out in accordance with the principles of animal care established by the EU Directive 2010/63/EU and in agreement with Spanish legislation (Royal Decree 53/2013), and were approved by the UPV/EHU Ethical Board of Animal Welfare (CEEA; reference M20_2020_014), as well as in compliance with ARRIVE guidelines (Percie du Sert et al., 2020).

### 2.2 Chronic unpredictable mild stress (CUMS) protocols

Three different CUMS protocols were implemented: CUMS 1, CUMS 2, and CUMS 3 (**Figure 1**). For CUMS 1 and CUMS 2 protocols, 12 mice were randomly assigned to the CUMS groups and 12 to the control groups, and all animals were group-housed. For CUMS 3 protocol, 16 mice were randomly assigned to the CUMS group and individually housed, and 16 to the control group, and were kept grouped. Animals from the control groups and their counterparts undergoing chronic stress were kept in separate rooms to avoid inadvertent stress exposure to controls (Willner, 2017a). The stress paradigms were designed based on previously described protocols, with some modifications, and differed in stress intensity and frequency, and protocol duration (Du Preez et al., 2021a; Elizalde et al., 2008; Erkizia-Santamaría et al., 2025b). In all cases, stressful stimuli were applied in a random manner and were not repeated in consecutive days.

**Figure 1.**
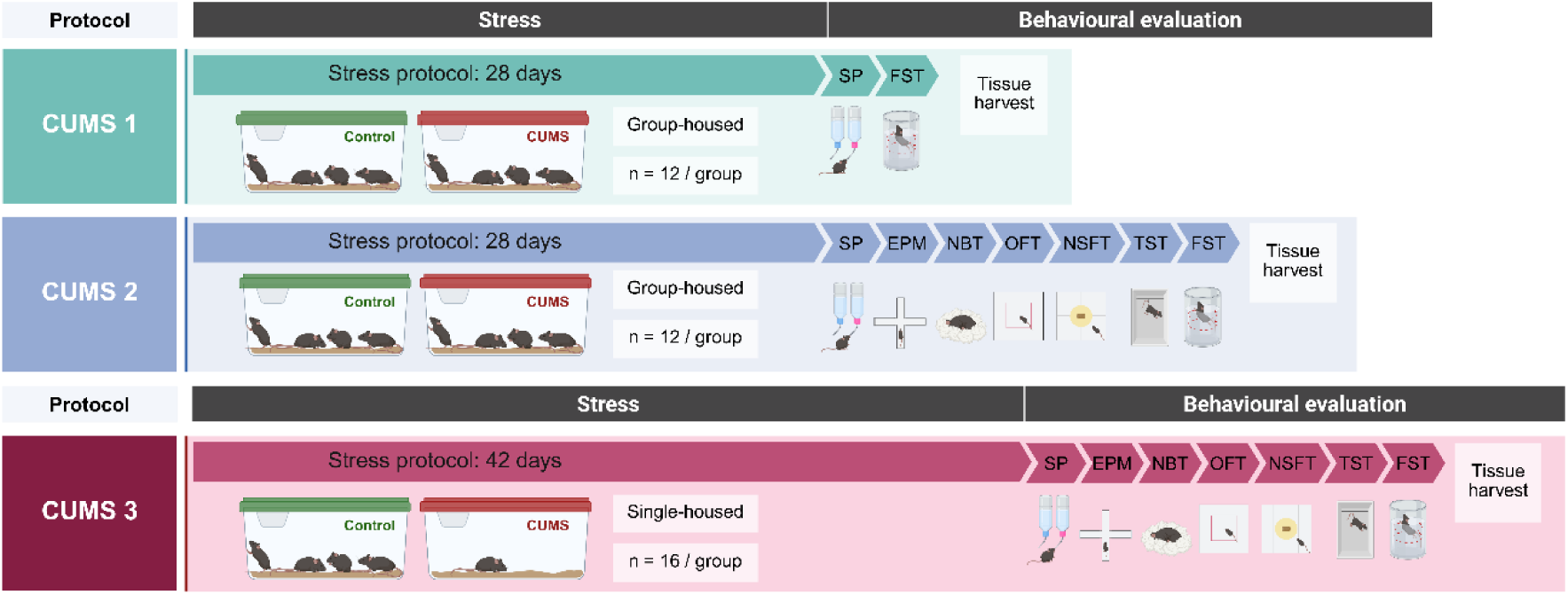
Timeline of CUMS 1, CUMS 2 and CUMS 3 stress paradigms, followed by behavioural evaluations and tissue harvest for physiological assessment. SP: sucrose preference test. FST: forced swimming test. EPM: elevated plus maze. NBT: nest-building test. OFT: open field test. NSFT: novelty-supressed feeding test. TST: tail suspension test.

CUMS 1 protocol had a duration of 28 days (4 weeks), and one single stressful stimulus was applied every day. CUMS 2 protocol also had a duration of 28 days (4 weeks), and three different stimuli were applied every day. CUMS 3 protocol had a duration of 42 days (6 weeks) and three to four different stimuli were applied every day. For CUMS 2 and CUMS 3 protocols, daily combinations of stressors were designed based on a point system to reach an equilibrium of stress intensity of 8-11 points every day (1: mild, 2: moderate, 3: high) (Erkizia-Santamaría et al., 2025b; Willner, 2017a) (**Table S1**).

### 2.3 Behavioural evaluations

Immediately following stress protocol cessation, animals were subjected to a battery of behavioural tests in order to evaluate the depressive- and anxiety-like phenotype of stressed animals induced by each CUMS paradigm. The order of assays was established according to recommendations, starting with the tasks requiring maximal effects of novelty and ending with the most stressful, as previously recommended (Du Preez et al., 2021a) (**Table S2**). All experiments were randomized and analysed in blind.

#### Sucrose preference test (SP)

SP was performed as previously described with minor modifications (Liu et al., 2018). Previous to the beginning of the stress protocol, a training phase was performed for 48 h in all groups of animals, in which bottle positions were switched every 24 h in order to avoid place preference. For the baseline measurement and test, an 8-h fasting period was set to promote water/sucrose (2%) consumption. Bottles were placed in cages at 17:00, and left for 15 h. In the CUMS 1 protocol, anhedonia was assessed in animals housed in groups of four per home cage, and the measured value represents the average consumption per group. In the case of CUMS 2 and 3, anhedonic state of all animals was evaluated individually, and afterwards control mice were returned to grouped home cages with food and water *ad libitum*. CUMS mice returned to the stress schedule as established. Sucrose preference was calculated as percentage ([sucrose consumption - water consumption] / [sucrose consumption + water consumption] x 100) (Cameron et al., 2023; Erkizia-Santamaría et al., 2025b). At baseline and testing phase for control animals, preference was considered above 65%, and data from mice that did not reach the criteria were removed for statistical analyses.

#### Elevated plus maze (EPM)

EPM was performed as previously described for evaluation of anxiety-like behaviour (Carobrez & Bertoglio, 2005). Time in the open arms in the duration of the test (5 min) was evaluated in the video tapes.

#### Nest-building test (NBT)

The spontaneous motivation of the animals was measured on the NBT. Each mouse was kept in an individual cage. One square of nesting material was introduced in each cage and mice were left undisturbed to build the nest. Nest-building skills were evaluated 30 min later. The nest quality was evaluated according to a scale using the following criteria: 1 (cotton square is intact), 2 (cotton square is partially used), 3 (cotton is scattered, but there is no form of nest), 4 (cotton is gathered but there is a flat nest), 5 (cotton is gathered to a ball-shaped nest) (Nollet, 2021).

#### Open field test (OFT)

The locomotor activity and emotionality from exploration was evaluated by the OFT (Seibenhener & Wooten, 2015). Mice were carefully placed in the centre of the arena and left to explore for 10 min (Gould et al., 2009). Videos were analysed using automated tracking software Smart 3.0 (PanLab SL, Barcelona, Spain). The total distance (cm) and time spent in the centre of the arena (s) were evaluated.

#### Novelty-suppressed feeding test (NSFT)

The NSFT was performed for the assessment of anxiety-like behaviour as previously described (Samuels & Hen, 2011). Food pellets were removed from cage grids 16 h prior to the NSFT. An open field arena was filled with sawdust and a single pellet attached to the ground was placed in the centre. The room was kept dark during the whole duration of the test except for a light bulb that illuminated the pellet (800–900 lux). Chewing or biting the pellet was established as the criterion to set the latency and remove the animal from the arena. A period of 10 min was set as the maximum latency time permitted.

#### Tail suspension test (TST)

Behavioural despair was evaluated using a TST apparatus (PanLab SL, Barcelona, Spain). Mice were hung from a piece of tape stuck to the tail. The duration of the test was 6 min (Steru et al., 1985), and the immobility time was manually evaluated by tape visualisation for the whole duration.

#### Forced swimming test (FST)

The modified Porsolt swim test was carried out as previously described (Lucki, 1997). A clear plastic cylinder (24 cm height × 20 cm diameter) was filled with water (24 ± 1 °C, 18 cm height). The mouse was determined to be immobile when floating in an upright position without other activity than the necessary to keep its head above water. Immobility was quantified during the last 4 min of the 6 min test, as previously described (Cryan et al., 2002).

### 2.4 Physiological evaluations

#### Bodyweight

All mice were weighed before the stress protocol, then randomly assigned to control or CUMS groups. Subsequently, mice were weighed weekly during chronic stress paradigm and before euthanasia. The bodyweight gain relative to baseline (%) was reported as a measure of a physical sign of chronic stress.

#### Food intake

Food intake was measured at baseline prior to the beginning of stress protocol, at the end of CUMS 2 protocol and CUMS 3 protocol. For all experimental groups, pellets in each cage were weighed at 17:00, and mice were left undisturbed to consume food and water overnight until 11:00 the following day. The amount of food consumed was corrected for bodyweight (g food/g bodyweight).

#### Tissue harvest

After behavioural evaluations, mice were euthanized through cervical dislocation. Peripheral tissues were dissected to obtain adrenal glands, spleen, brown adipose tissue (BAT), and white adipose tissue (WAT). Tissues were weighed immediately after dissection.

### 2.5 Statistical analysis

Normality of data distribution was assessed using the Shapiro–Wilk test prior to subsequent statistical analyses. Body weight changes over time during the CUMS protocols were analyzed using two-way repeated-measures ANOVA, followed by Bonferroni post hoc tests when a statistically significant interaction was detected. Factors were identified as F_t_ (time), F_CUMS_ (CUMS), and F_i_ (interaction). For statistical analysis of physiological and behavioural data, unpaired *t-*test was used (control vs stressed). For analysis of scores in NBT (semi quantitative parameter, considered categorical ordinal data), non-parametric Mann-Whitney test was performed (control vs stressed). Statistical outliers were identified using Grubbs’ test. All results are shown as mean±SEM. In all cases, statistical significance was considered when p<0.05. Data were analysed using GraphPad Prism™ software version 10.5 (GraphPad Software Inc. CA, USA).

## 3. RESULTS

### 3.1 Effect of chronic stress paradigms on physiological measures

Firstly, physiological parameters known to be altered in consequence of chronic stress were investigated following CUMS 1, CUMS 2 and CUMS 3 protocols. Changes in body weight throughout the chronic stress protocols were analyzed using two-way repeated-measures ANOVA. In the CUMS 1 paradigm, a significant main effect of time was observed, whereas neither the effect of chronic stress nor the time x stress interaction reached statistical significance (F_t_(4,88)=91.28, p<0.0001; F_CUMS_(1,22)=0.06, p=0.81; F_i_(4,88)=2.18 p=0.08) (**Figure 2a**). Bodyweight gain relative to baseline was not different between control and CUMS 1 groups (t=1.89, p=0.08) (**Figure 2b**). In CUMS 2, a significant effect of time and a significant interaction between factors, but no effect of stress was revealed (F_t_(1.98,43.60)=73.95, p<0.0001; F_CUMS_(1,22)=0.63, p=0.43; F_i_(4,88)=15.42, p<0.0001) (**Figure 2c**). Moreover, bodyweight gain was significantly different between control and CUMS 2 groups (t=2.44, p<0.05) (**Figure 2d**). Finally, in CUMS 3 protocol, a significant effect of time, stress and interaction between factors was found (F_t_(2.83,84.81)=360.60, p<0.0001; F_CUMS_(1,30)=22.63, p<0.0001; F_i_(6,180)=34.37, p<0.0001). Additionally, Bonferroni *post hoc* test revealed differences in bodyweight in weeks 3, 4, 5, and 6 between control and stressed groups (week 3 t=3.34, p<0.05; week 4 t=4.10, p<0.01; week 5 t=5.80, p<0.0001; week 6 t=10.46, p<0.0001) (**Figure 2e**). Expectedly, bodyweight gain was significantly different between control and CUMS 3 groups (t=8.46, p<0.0001) (**Figure 2f**).

**Figure 2.**
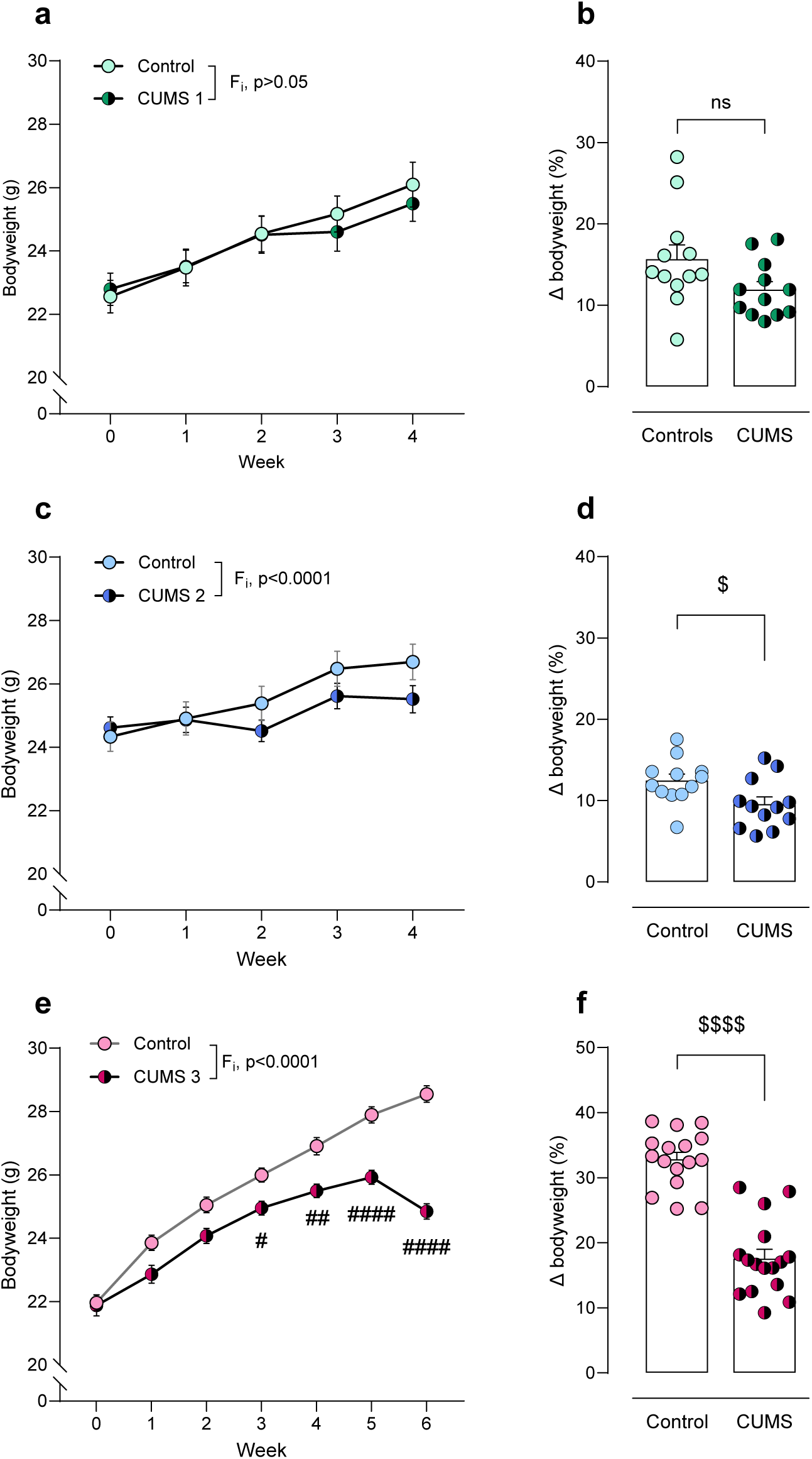
Bodyweight of control and stressed mice during CUMS 1 (**a, b**), CUMS 2 (**c, d**), and CUMS 3 (**e, f**) protocols. Two-way repeated measures ANOVA followed by Bonferroni *post hoc* test. ^#^p<0.05, ^##^p<0.01, ^####^p<0.0001. Unpaired *t*-test. ^$^p<0.05, ^$$$$^p<0.0001. ns, non significant, p>0.05.

The effects of chronic stress exerted by the CUMS 1 protocol were considered insufficient to induce a robust depression-like model, as evidenced by the absence of impact on the animals’ weight gain (**Figure 2b**) and the absence of alterations in the depression-like phenotype (**Figure S1**), in spite of mild adrenal gland hypertrophy. Therefore, a complete assessment of the physiological and behavioural changes in consequence of chronic stress was carried out in CUMS 2 and CUMS 3 protocols.

Food intake was verified to be statistically no different between groups prior to stress (data not shown). Following stress protocol, food consumption was measured again and normalized by bodyweight. Food intake was significantly reduced in stressed animals under the CUMS 2 protocol (t=3.03, p<0.01), whereas it was significantly increased in those subjected to CUMS 3 (t=8.22, p<0.0001) (**Figure 3a**). A non-significant increase in adrenal gland weight was observed in CUMS 2 stressed group (t=1.46, p=0.16) (**Figure 3b**). Moreover, spleen weight and BAT weight were not statistically different between control and CUMS 2 stressed group (t=0.94, p=0.36; t=2.06, p=0.05, respectively) (**Figure 3c, d**). Nonetheless, CUMS 3 stressed group showed a significant increase in adrenal gland weight (t=11.76, p<0.0001), a significant decrease in spleen weight (t=9.70, p<0.0001), and a significant increase in BAT weight (t=3.45, p<0.01) compared to controls (**Figure 3b, c, d**). Regarding WAT, a significant decrease was found in stressed mice in CUMS 2 (t=2.17, p<0.05) and CUMS 3 (t=3.58, p<0.01) compared to their respective controls (**Figure 3e**), consistent with the previously observed attenuation in body weight gain.

**Figure 3.**
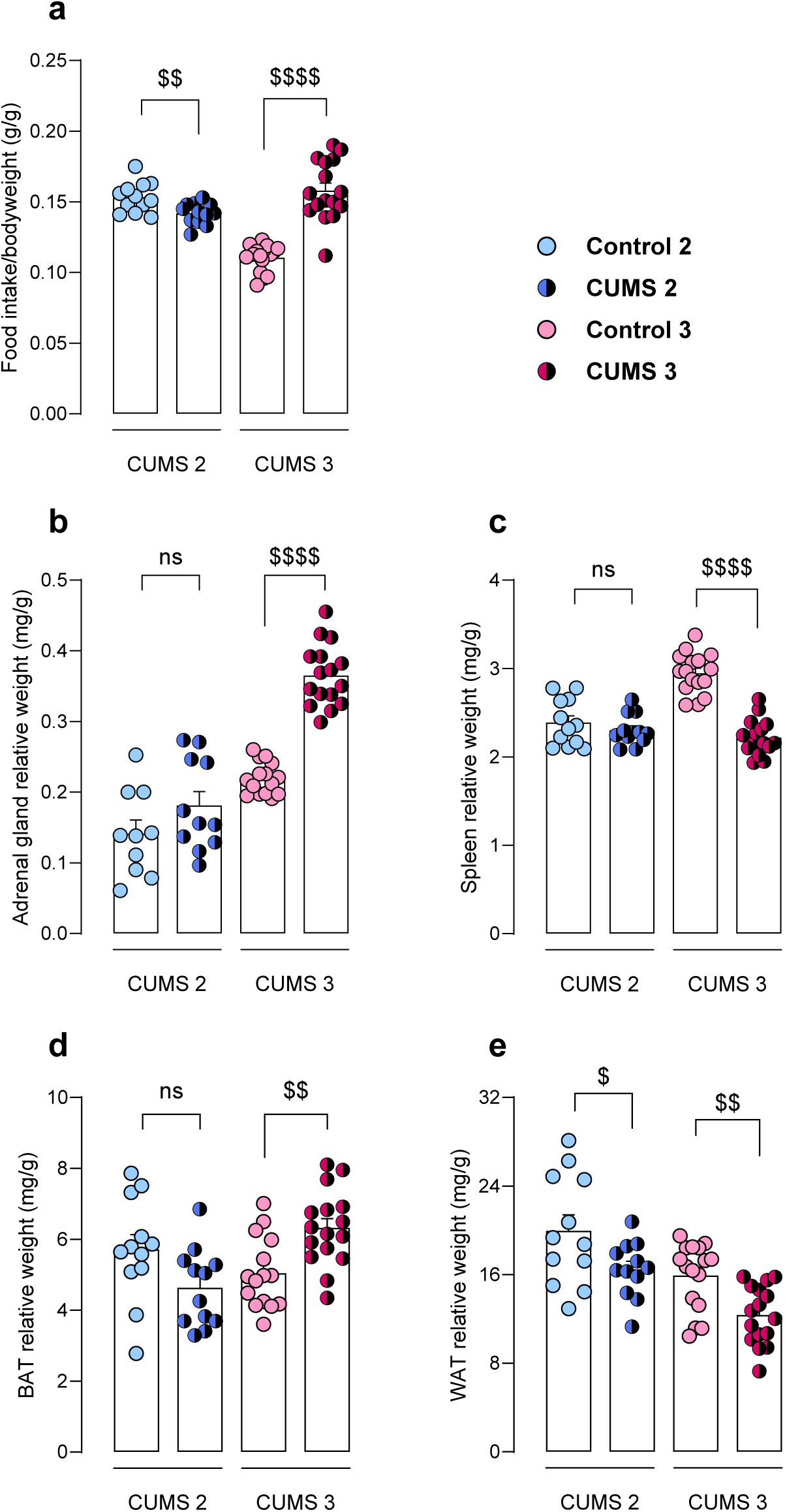
Physiological evaluation of animals in CUMS 2 (n=12) and CUMS 3 (n=16) protocols. Food intake (**a**), adrenal gland weight (**b**), spleen weight (**c**), white adipose tissue (WAT) weight (**d**) and brown adipose tissue (**e**) relative to bodyweight. Unpaired *t*-test. ^$^p<0.05, ^$$^p<0.01, ^$$$$^p<0.0001. ns, non significant, p>0.05.

### 3.2 Effect of chronic stress paradigms on depression-like phenotype

Anhedonic state was assessed by evaluation of SP. Preference was verified to be statistically no different between groups at baseline (data not shown). Stressed animals in CUMS 2 did not exhibit a significant difference in preference compared to controls (t=1.20, p=0.25). On the contrary, stressed animals in CUMS 3 showed significantly lower preference relative to controls (t=3.10, p<0.01) (**Figure 4a**). Spontaneous motivation was assessed by NBT. In CUMS 2 paradigm, stressed mice did not show a significant difference in NBT compared to controls (Mann-Whitney U=66.50, p=0.90), while stressed mice in CUMS 3 showed a significant decrease in nest-building score (Mann-Whitney U=3, p<0.0001) (**Figure 4b**).

**Figure 4.**
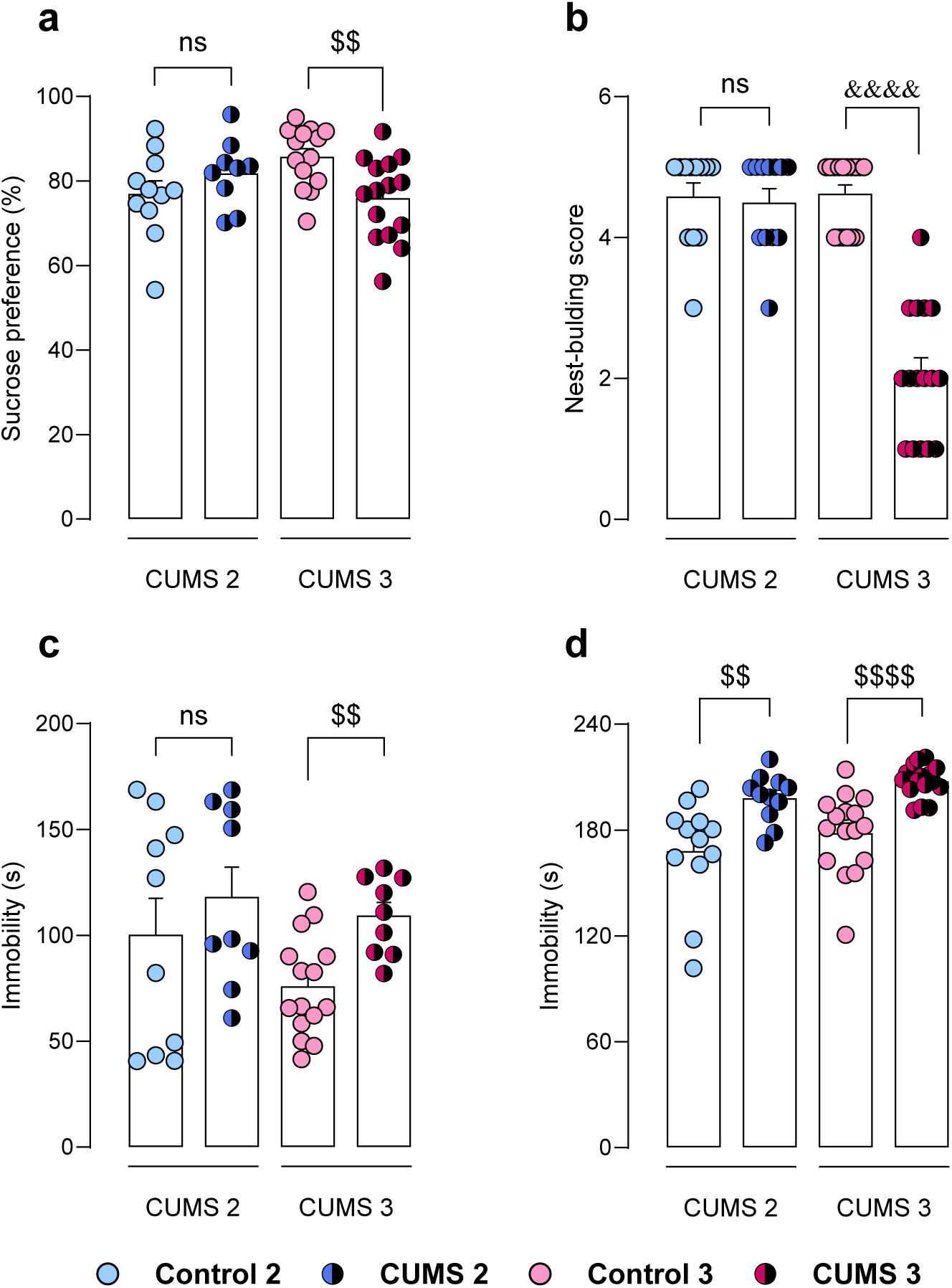
Evaluation of depressive-like phenotype of animals in CUMS 2 (n=12) and CUMS 3 (n=16) protocols. Sucrose preference (**a**), nest-building score in NBT (**b**), TST immobility time (**c**) and FST immobility time (**d**). Unpaired *t*-test. ^$$^p<0.01, ^$$$$^p<0.0001. Mann-Whitney test. ^&&&&^p<0.0001. ns, non significant, p>0.05.

Behavioural despair was assessed by TST and FST, where immobility-times were evaluated. In TST, stressed mice in CUMS 2 protocol did not exhibit significant differences in immobility compared to controls (t=0.81, p=0.43), whereas stressed group in CUMS 3 showed significantly higher immobility-time relative to controls (t=3.84, p<0.01) (**Figure 4c**). In FST, stressed groups from both CUMS 2 and CUMS 3 protocols exhibited significantly higher immobility times compared to their corresponding control counterparts (t=3.13, p<0.01; t=4.80, p<0.001, respectively) (**Figure 4d**).

### 3.3 Effect of chronic stress paradigms on anxiety-like phenotype

Anxiety-like phenotype induced by chronic stress paradigms was evaluated in several behavioural tests. Firstly, latency to food in NSFT was not statistically different between stressed mice in CUMS 2 and control group (t=0.43, p=0.67). On the contrary, stressed mice in CUMS 3 showed significantly higher latency to food compared to control counterparts (t=2.17, p<0.05) (**Figure 5a**). Similarly, in EPM, stressed mice in CUMS 2 protocol did not exhibit significant differences in time spent in open arms compared to controls (t=0.17, p=0.87), whereas stressed mice in CUMS 3 showed significantly lower time in open arms (t=2.37, p<0.05) (**Figure 5b**).

**Figure 5.**
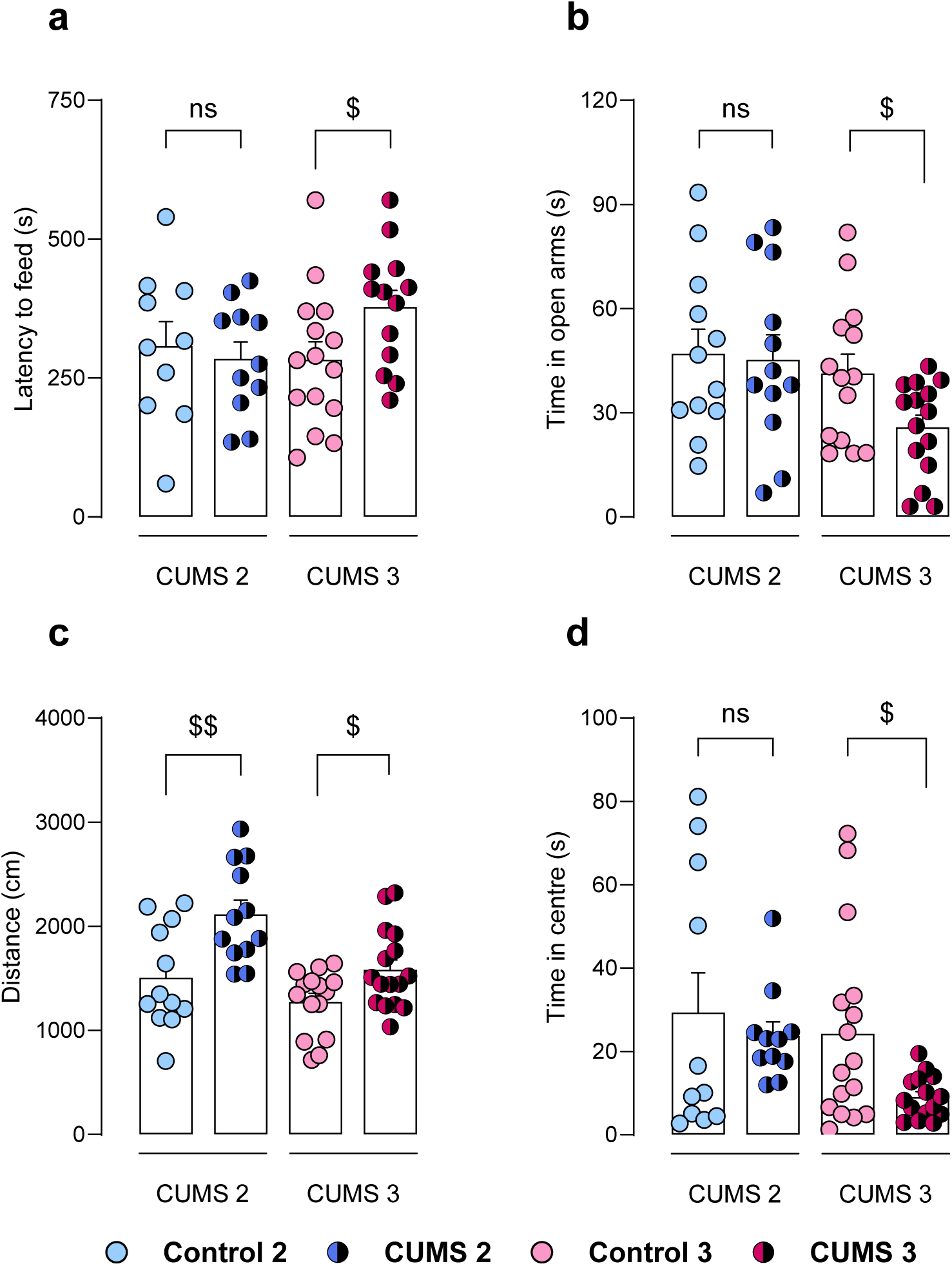
Evaluation of anxiety-like phenotype of animals in CUMS 2 (n=12) and CUMS 3 (n=16) protocols. Latency to feed in NSFT (**a**), time in open arms in EPM (**b**), travelled distance (**c**) and time in centre (**d**) in OFT. Unpaired *t*-test. ^$^p<0.05, ^$$^p<0.01. ns, non significant, p>0.05.

Exploratory activity and anxiety-like phenotype were also evaluated in OFT. A significant increase in travelled distance was exhibited by stressed animals in comparison to controls, both in CUMS 2 (t=3.09, p<0.01) and CUMS 3 (t=2.48, p<0.05) protocols (**Figure 5c**). Regarding time in the centre of the arena, no significant differences were found between stressed mice in CUMS 2 and controls (t=0.55, p=0.59). Meanwhile, stressed animals in CUMS 3 spent significantly less time in the centre of the OFT (t=2.62, p<0.05) (**Figure 5d**).

## 4. DISCUSSION

Preclinical models stand at the core of translational research, serving as valuable tools to investigate the neurobiological mechanisms underlying human brain disorders. Despite the shortcomings and limitations of animal models of disease, their use has significantly contributed to the elucidation of pathophysiological mechanisms of depressive disorders, and to the rational identification of novel molecules with antidepressant action (Bale et al., 2019; Kalueff et al., 2007). Chronic stress based rodent models in particular are considered to have a greater aetiological relevance and face validity than other depression-like animal models (Strekalova et al., 2011). Notwithstanding, the widely used CUMS model has been perceived as insufficiently reliable and reproducible: many research laboratories have experienced difficulties establishing or replicating the procedures (Willner, 2017a). Furthermore, some have reported anomalous or paradoxical effects of CUMS, such as increased sucrose intake, decreased scores of helplessness and anxiolytic-like features, which yield discordance between the phenotype of chronically stressed rodents and human symptoms of depression (Strekalova et al., 2022). In the present work, we developed different CUMS paradigms with the aim to identify experimental variables that may contribute to the effective induction of depression-related behavioural and physiological impairments in mice within a consistent laboratory setting.

Reduced bodyweight or weight gain in CUMS-exposed rodents has been considered an indicator of sufficient stress load that could lead to the induction of anhedonia (Strekalova et al., 2022). In the present work, no effects were observed on bodyweight changes during or following CUMS 1 protocol, a slight decrease in weight gain was detected in stressed mice following CUMS 2, and a robust reduction in weight gain was observed in stressed mice in CUMS 3. Such disparities may be attributable to the progressive increase in stress intensity and frequency of each paradigm. Evidence from the literature seems to suggest that significant differences in bodyweight between control and stressed mice from the C57BL/6 strain emerge in an unsteady time-interval, starting as soon as week 2 of stress exposure (Huang et al., 2018; Li et al., 2018; Monteiro et al., 2015) and as late as week 4 (Pałucha-Poniewiera et al., 2020). In other cases, no differences arise at all, even in the presence of a depressive-like behavioural phenotype (Elizalde et al., 2008; Pothion et al., 2004). In the current work, significant differences in bodyweight between controls and stressed mice in CUMS 3 protocol arise within 3 weeks of stress onset. Such changes appear to be a sign of potential behavioural impairments of depressive nature, and are often an accompanying feature following chronic stress protocol induction, prompting its use as a standard for adjusting stress intensity (Strekalova et al., 2022). However, in accordance with several reports, associations between changes in bodyweight and depressive-like phenotype are not warranted (Strekalova & Steinbusch, 2010). Therefore, loss of weight should not be considered an unequivocal characteristic of the effects of chronic stress in mice. While it may serve as an easily measurable initial indicator that the model is progressing toward the desired disease phenotype, it should not be relied upon as the sole criterion for justifying investigations into the neurobiology of depression or anxiety.

Chronically stressed mice from CUMS 3 protocol exhibited several macroscopic physiological alterations, i.e. marked adrenal hypertrophy, decreased WAT weight and increased food intake. In relation to HPA hyperactivity, evidenced by the hypertrophy of adrenal glands, continuous glucocorticoid release from the adrenal cortex enhances catabolic metabolism during stress (Michel et al., 2005). This phenomenon has been linked to increased energy expenditure and loss of body fat and lean body mass in mice subjected to chronic variable stress (Kuti et al., 2022). Moreover, chronic stress exposure has been seen to induce increased food intake accompanied by decreased bodyweight gain in rats (Abulmeaty et al., 2023), in accordance with the present results from the CUMS 3 protocol. Notably, evidence from human and preclinical studies has shown that stress may induce overeating, but can also lead to decreased caloric consumption (Francois et al., 2022). The present results also suggest that chronic stress may result in conflicting effects in the feeding of mice, as stressed animals in the CUMS 2 protocol exhibited decreased food intake, whereas those subjected to the CUMS 3 protocol showed a significant increase in food consumption compared to controls. Individual housing in CUMS 3 stressed mice may have been a determinant factor for such increased feeding, as previous reports have demonstrated that social isolation is associated with higher food intake in mice (Benfato et al., 2022; Yamada et al., 2015). This contrasting pattern in grouped vs isolated mice may reflect a shift toward an anxious or impulsive phenotype under prolonged isolation stress.

It is known that high adrenocortical stimulation is associated with the breakdown of fat storages in the body (Clemente-Suárez et al., 2023), in accordance with the present data from CUMS 2 and CUMS 3 protocols. WAT, predominant site for energy storage, is an active endocrine tissue, which produces adipokines, signalling molecules that regulate a host of processes, e.g. appetite, fat distribution, energy expenditure, mood and cognitive performance (Fasshauer & Blüher, 2015; Lee et al., 2019; Wang et al., 2023). Importantly, dysregulation of adipokines has been implicated in the neurobiology of depressive disorders, and proposed as a potential therapeutic target (Formolo et al., 2019). For instance, the appetite-suppressing hormone leptin has been found to be decreased in non-obese patients with depression (Morris et al., 2012), and has been suggested to be implicated in the pathophysiology of the disease (Clemente-Suárez et al., 2023; Yang et al., 2016). In addition, BAT is the main non-shivering thermogenic site in mammals and plays a crucial role in heat production and energy balance in rodents (Fernandes et al., 2023; Sun et al., 2014). In the present work, chronic stress-exposed mice in CUMS 3 protocol exhibited increased relative BAT weight. Previous works have established that social isolation may give rise to an enlarged BAT development, as well as adaptations in its function to maintain core body temperature (Fernandes et al., 2023; Sun et al., 2014). Additionally, stressed mice in CUMS 3 protocol exhibited decreased spleen weight. Other works have described atrophic decreases in spleen weight, possibly linked to stress-activated immune response, even though the precise pathological mechanisms are not entirely clear (Sarjan & Yajurvedi, 2019; Smaniotto et al., 2023).

The current report portrays a complete and comprehensive characterization of the phenotypic alterations induced by chronic stress in the protocols CUMS 2 and CUMS 3. We executed a wide battery of behavioural tests, since the evaluation of each symptomatic domain through multiple tests enhances the reliability and consistency of results, and improves their translational relevance to human contexts (Erkizia-Santamaria et al., 2025a). The SP paradigm is the most frequently used method for assessing anhedonia in rodent models, as it measures hedonic sensitivity rather than reward-seeking behaviour (Strekalova et al., 2022). CUMS rodent models have been found to be strongly associated with anhedonic behaviour, even though high heterogeneity has been observed across studies, likely related to factors like the nature of stressors, housing conditions and strain (Antoniuk et al., 2019; Strekalova et al., 2022). In line with such heterogeneity, in the present work, only stressed mice from CUMS 3 protocol exhibited anhedonia in the SP. Additionally, spontaneous motivation and behavioural despair were also impaired in stressed mice in CUMS 3 protocol. Likewise, we observed enhanced anxiety-like behaviours in consequence of CUMS 3 protocol in a multiple assays. Thus, the CUMS 3 paradigm recapitulates an archetypical disease-like phenotypic profile of chronic stress-based depression-like models (see **Table S3** for a summary of the physiological and behavioural findings in the current study).

The phenotypic discrepancies observed across the different stress protocols here presented may be accounted for by two key factors. On the one hand, the increased stress intensity and protocol duration of CUMS 3 is likely a contributing element to the overall behavioural and physiological impairments detected. Some works in the literature support the present findings. For instance, alterations indicative of behavioural despair and anxious phenotype were only evident after 8 weeks of stress, but not 4 weeks, in an elegant comparative study in mice (Monteiro et al., 2015). The relationship between chronic stress and psychopathology is closely tied to factors related to stress intensity and duration. Individuals can successfully adapt to stress across a wide range of intensity and span, while undergoing physiological and behavioural changes to adjust to prolonged or intermittent stress exposure (**Figure 6**). In fact, relatively low to moderate stress can enhance cognitive processes such as memory consolidation, processing and encoding (Brosens et al., 2024), improving the organism’s performance. The ability or inability of an organism to elicit adaptive responses-that is, contextually appropriate physiological or behavioural reactions-is likely influenced by several factors contributing to stress susceptibility, including genetic predisposition, sex, and age, among others. The ensemble of these factors ultimately dictates the amount of stress exposure that can be tolerated, and is directly conditioned by the intensity and duration of it (Radley & Herman, 2023). In relation to the present data, a comparison between behavioural and physiological outcomes from the different CUMS protocols described in this report sheds light onto the shift from adaptation to pathology, that is, how the organism may undergo adaptive responses in an adjustment phase of stress reactivity, or rather develop impairments associated to pathology.

**Figure 6.**
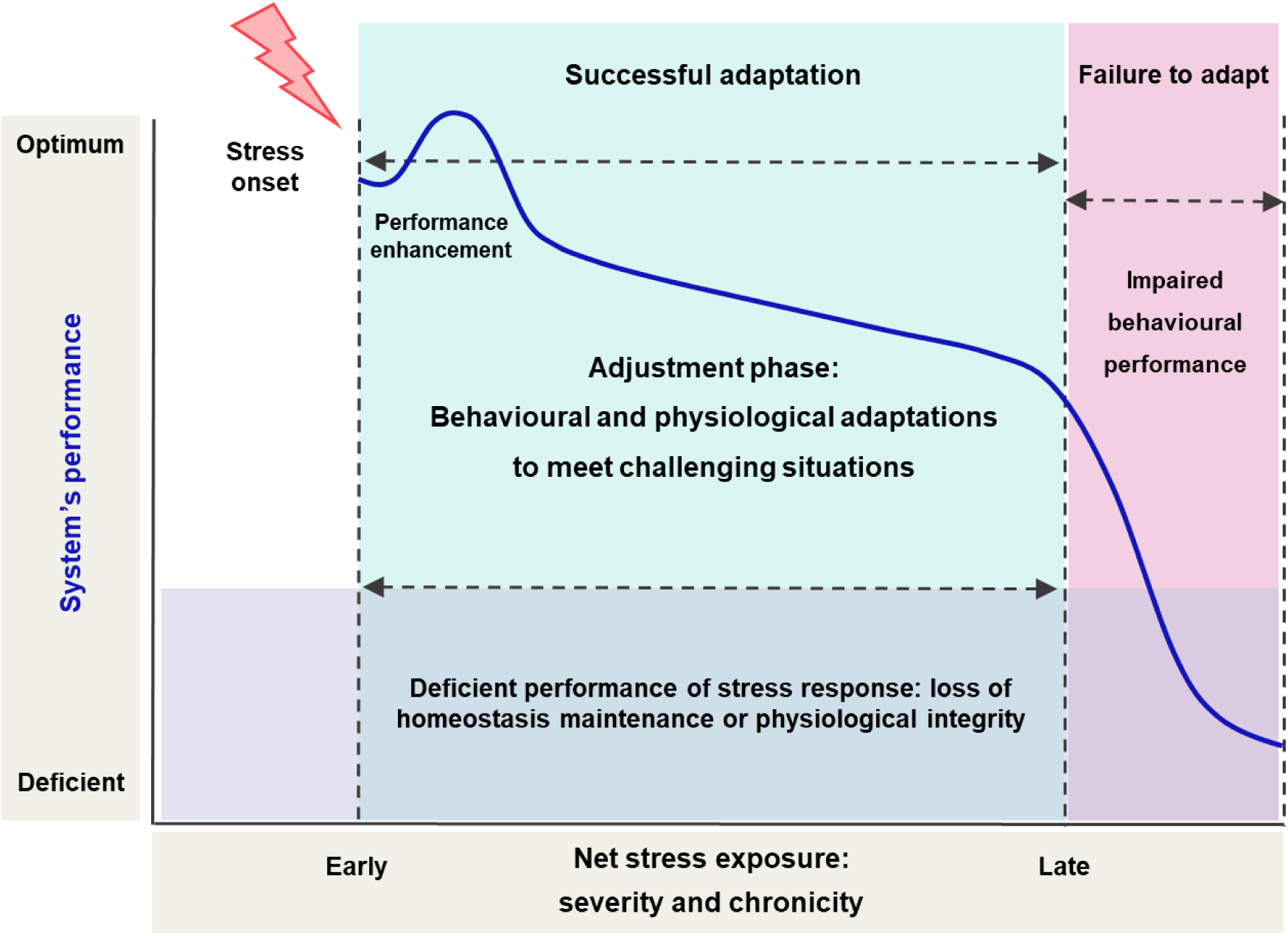
Graphical illustration of the path from adaptation to pathology in progression of chronic stress. The blue line represents the organism’s progression in response to stress. Early stress may enhance processes like memory strength and precision, to facilitate survival. From stress onset, the organism initiates adaptive changes that involve physiological and behavioural adjustments to meet the demands of the stressful situation -conservative behavioural choices to limit exposure to danger and energy expenditure-(blue shaded area). Prolonged or intense stress may degrade the system’s performance and result in adaptation failure -inappropriate behavioural or physiological response-(purple and pink shaded areas). Adapted from Radley & Herman, 2023.

On the other hand, social isolation was introduced in CUMS 3 protocol as a permanent stressful stimulus. Importantly, social isolation has been associated with higher risk of mental health problems, including depression (Cacioppo et al., 2006; Chou et al., 2011), whereas social buffering may have positive effects on health and stress responses in relation to the neuroendocrine system and to behaviour (Du Preez et al., 2021b; Kikusui et al., 2006; Muscat & Willner, 1992). The present results reinforce the argument that social isolation may be a strong precipitant for behavioural and neuroendocrine impairments in rodents. In the experiments conducted in group-housed animals, absent or weak disease-relevant phenotypes were observed in stress-exposed mice. These results are in disagreement with a myriad of publications that have reported robust behavioural, neurochemical and biochemical alterations induced by stress protocols with comparable features to CUMS 1 or CUMS 2, i.e. grouped housing, 4 weeks or less of duration, low-medium frequency of stressful stimuli (Al-Hasani et al., 2013; Burstein & Doron, 2018; Jung et al., 2014; Pałucha-Poniewiera et al., 2020). Meanwhile, stressed mice in CUMS 3 protocols exhibited robust impairments in physiological and behavioural parameters associated to depressive disorders. Collectively, these data suggest that longer and more intense chronic stress protocols in socially isolated C57BL/6J mice enhance the induction of disease-relevant deficits.

It is important to acknowledge certain limitations of the present study. Firstly, all experiments were conducted exclusively in male mice, as is the case for the majority of preclinical research on chronic stress (Du Preez et al., 2021b). The sex bias in the literature of preclinical studies of chronic stress-based models represents a major limitation, particularly since the prevalence of depression is significantly higher in women (Albert, 2015; Malhi & Mann, 2018), and clearly reinforces that research using female rodents is indeed underrepresented and necessary (Beery & Zucker, 2011; Prendergast et al., 2014). It is also worth empathizing that only a few key factors contributing to stress-induced neurobiological effects have been modified and tested in the present work, i.e. duration and frequency of stress exposure and housing conditions. The concurrent modifications in such factors between protocols prevents us from discerning the weight of each individual variable contributing to the behavioural and physiological consequences of chronic stress. In view of the divergences identified between CUMS protocols, it remains plausible that controllable (stress duration, intensity, strain, etc.) and unknown environmental or methodological factors could differentially alter behavioural and pathophysiological outcomes when testing animal models of disease. Indeed, a significant source of unreliability may be associated with the inadvertent stress experienced by control animals. This could easily occur when control and CUMS mice are accommodated in the same room, which should be strictly avoided, as was in the present work (Willner, 2017a). Lastly, publication bias has been reported to significantly undermine the reliability of CUMS studies. On the one hand, bias may emerge from the poor description of the CUMS method or lack of details in housing condition of animals, giving rise to high levels of uncertainty (Lages et al., 2021). On the other hand, it is likely that studies with non-significant findings may have been left unreported. Therefore, the published data could represent the tip of an iceberg with evidence of unreliability below the surface, and truthfulness of data may be compromised by a large amount of unpublished records (Du Preez et al., 2021b; Markov, 2022; Willner, 2017a).

## 5. CONCLUSIONS

Monitoring body weight remains a key primary indicator for evaluating the potential impact of a chronic stress regimen. To corroborate the physiological consequences of stress exposure, adrenal gland hypertrophy serves as a reliable marker of HPA axis activation. However, defining a pathological phenotype requires validation through a comprehensive behavioural test battery, ideally including multiple assays per symptom domain to reduce the risk of false positives or negatives. This multidimensional approach allows for a more accurate assessment of the model’s position along the pathological trajectory and enhances its translational value. Notably, differences in individual susceptibility—as influenced by factors such as mouse strain or substrain—as well as variations in stressor intensity, type, and housing conditions can critically affect behavioural outcomes. These factors, along with protocol-related variability across laboratories, likely underlie many of the discrepancies reported in the literature. Therefore, detailed and transparent reporting of all protocol parameters is essential to improve reproducibility and facilitate meaningful comparisons across studies. When applied to well-characterized models, this level of methodological rigour significantly strengthens the reliability of pharmacological screening and increases the likelihood of successful clinical translation.

## CRediT authorship contribution statement

**Ines Erkizia-Santamaría**: Conceptualization, Formal analysis, Investigation, Methodology, Writing - original draft. **Luna Fernández**: Formal analysis. **Igor Horrillo:** Conceptualization, Methodology, **J. Javier Meana**: Resources, Supervision, Writing – review and editing. **Jorge E. Ortega:** Conceptualization, Methodology, Resources, Supervision, Writing – review and editing.

