## Supplemental Figure S1, Table S1-S3 for "Behavioural and Physiological Signatures of Chronic Unpredictable Mild Stress Models in Mice"

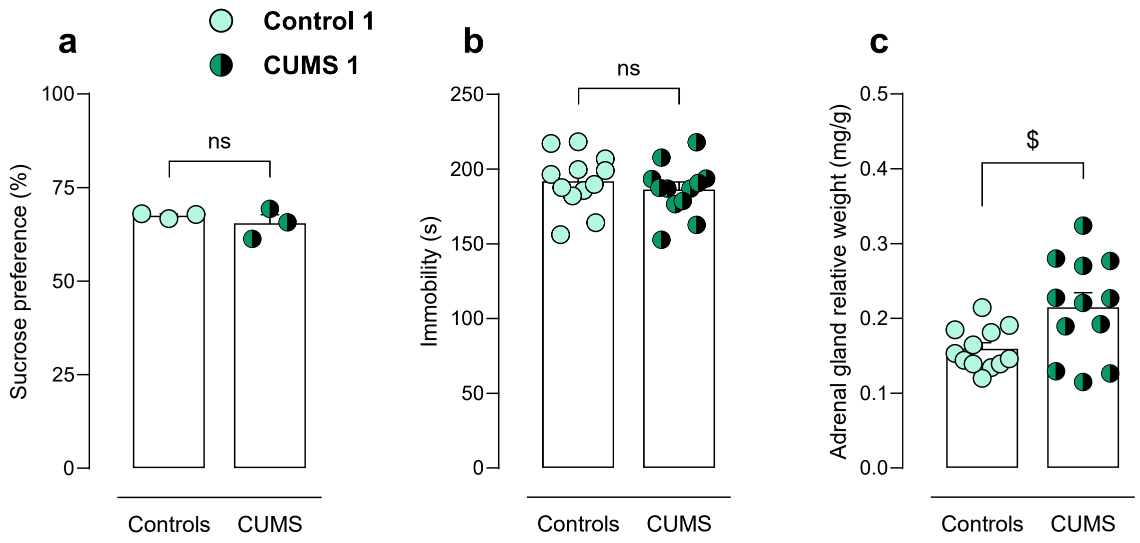


**Figure S1.** Behavioural and physiological evaluation of stressed and control animals in CUMS 1 protocol (n=12). **a**. Sucrose preference (%) (grouped analysis, n=3 cages/group). **b**. Immobility time in FST. **c.** Adrenal gland weight relative to bodyweight (mg/g). Unpaired *t*-test. $^{\$}p<0.05$. ns, non significant, p>0.05.

**Table S1.** Stressful stimuli and description for each CUMS protocol: stressor, details, duration, phase of the cycle (D: dark, active; L: light, inactive phase) and intensity according to the severity point system. *Food/water deprivation for CUMS 3 protocol was applied for 16 hours, only before SP (food and water) and NSFT (only food) tests.

| *Stressor* | *Description* | *Duration* | *Phase of the cycle* | *Intensity* | *1* | *2* | *3* |
|---|---|---|---|---|---|---|---|
| **Cage tilting** | Cages were tilted sideways 45º | 12 h | D | 1 | ✓ | ✓ | ✓ |
| **Food/water deprivation** | Food and water were removed | 8 h / 16 h | D | 1 | ✓ | ✓ | ✓ * |
| **Space reduction** | A clear plastic wall was placed in the cage to reduce habitable space to 1/3 | 12 h | D | 1 | | ✓ | |
| **Cage swap** | Sawdust from cages within the stress group was swapped | - | L | 1 | | ✓ | |
| **Overcrowding** | 8 animals were placed in an individual cage (size) | 2 h | L | 1 | | | ✓ |
| **Alarm clock** | An alarm clock at 85 dB was activated at random times | 10 min | L + D | 2 | ✓ | ✓ | ✓ |
| **White noise** | An untuned radio was set at 85 dB | 4 h | L | 2 | ✓ | ✓ | ✓ |
| **Wet bedding** | 200 mL of water were poured over 400 mL of sawdust | 12 h | D | 2 | ✓ | ✓ | ✓ |
| **No bedding** | Sawdust was removed | 12 h | D | 2 | ✓ | ✓ | ✓ |
| **Continuous light** | Lights were kept on for on entire sleep cycle | 24 h | L + D | 2 | ✓ | ✓ | ✓ |
| **Light pulses** | Lights were switched on and off every 20 min | 12 h | D | 2 | | ✓ | ✓ |
| **Predator odour** | Sawdust was removed and replaced with sawdust from rats' cages | 2 h | L | 2 | | ✓ | ✓ |
| **Cold exposure** | Cages were introduced in a chamber at 4 ºC | 1 h | L | 3 | ✓ | ✓ | ✓ |
| **Heat exposure** | A heat source was placed in the rack. Temperature did not exceed 40 ºC | 2 h | L | 3 | | ✓ | ✓ |
| **Stroboscopic lights** | Flashing lights were applied | 4 h | L | 3 | | ✓ | ✓ |
| **Water bath** | Sawdust was removed and replaced with warm (23 ºC) water (2 cm-high) | 2 h | L | 3 | | ✓ | ✓ |
| **Restraint (a)** | Mice were introduced in ventilated plastic cups (7 x 5 cm) | 2 h | L | 3 | | ✓ | |
| **Restraint (b)** | Mice were introduced in ventilated plastic falcons (12 x 3 cm) | 2 h | L | 3 | | | ✓ |

**Table S2**. Duration of chronic stress protocol and days of behavioural testing in CUMS 1, CUMS 2 and CUMS 3 paradigms.

| | | **CUMS 1** | **CUMS 2** | **CUMS 3** |
|---|---|---|---|---|
| **Behavioural test** | **Protocol duration** | 28 days | 28 days | 42 days |
| Sucrose Preference test (SP) | | Day 28 | Day 28 | Day 42 |
| Elevated Plus Maze (EPM) | | - | Day 30 | Day 43 |
| Nest-Building test (NBT) | | **-** | Day 31 | Day 44 |
| Open Field test (OFT) | | **-** | Day 32 | Day 45 |
| Novelty-Suppressed Feeding test (NSFT) | | - | Day 33 | Day 46 |
| Tail-Suspension test (TST) | | - | Day 34 | Day 47 |
| Forced Swimming test (FST) | | Day 30 | Day 35 | Day 48 |
| Tissue harvest | | Day 32 | Day 36 | Day 49 |

**Table S3**. Summary of physiological and behavioural readouts in stressed groups from CUMS 1, CUMS 2 and CUMS 3 protocols. ↑, increase**.** ↓, decrease. =, no change.

| | ***CUMS protocol*** | | |
|---|---|---|---|
| ***Parameter*** | ***1*** | ***2*** | ***3*** |
| Bodyweight gain (%) | = | ↓ | ↓ |
| Food intake (g/g) | | ↓ | ↑ |
| Adrenal gland weight (mg/g) | ↑ | = | ↑ |
| Spleen weight (mg/g) | | = | ↓ |
| BAT weight (mg/g) | | = | ↑ |
| WAT weight (mg/g) | | ↓ | ↓ |
| Sucrose preference (%) | = | = | ↓ |
| Nest-building score | | = | ↓ |
| TST immobility time (s) | | = | ↑ |
| FST immobility time (s) | = | ↑ | ↑ |
| NSFT latency to feed (s) | | = | ↑ |
| EPM time in open arms (s) | | = | ↓ |
| OFT distance (cm) | | ↑ | ↑ |
| OFT time in centre (s) | | = | ↓ |
